# Endoreduplication in *Drosophila melanogaster* progeny after exposure to acute γ-irradiation

**DOI:** 10.1101/376145

**Authors:** Daria A. Skorobagatko, Alexey A. Mazilov, Volodymyr Yu. Strashnyuk

## Abstract

The purpose of investigation was to study the effect of acute γ-irradiation of parent adults on the endoreduplication in *Drosophila melanogaster* progeny. As a material used *Oregon-R* strain. Virgin females and males of *Drosophila* adults at the age of 3 days were exposed at the doses of 8, 16 and 25 Gy. Giant chromosomes were studied at late 3rd instar larvae by cytomorphometric method. The average polyteny in males increased at the dose of 8 Gy by 10.6%, at the dose of 25 Gy by 7.4%, and did not change at the dose of 16 Gy. In females, the polyteny did not differ from the control value irrespective of the irradiation dose. The statistical impact power of sex on polyteny was 4,9%, the radiation impact was 26,8%. The enhancement of endoruplication is considered as a consequence of an increasing selection pressure after irradiation.

## Introduction

It is known that ionizing radiation can influence the genetic apparatus of cells, causing damage to DNA and mutations (Dubrova 2006, Golub and Chernyk 2008, Vasil’eva et al. 2011, Skorobogatko et al. 2015). This is due to both direct and indirect effects of irradiation.

Ionizing radiation also affects the cell cycle, the passage of its individual stages. A well-known effect is a sharp decrease in the mitotic index, the so-called “radiation block of mitoses” (Deckbar et al. 2011). In addition, the delay G_1_/S-transition (G_1_-block) and the transition from G_2_ phase to M phase (G_2_-block) is possible. Sometimes there are opposite effects: an increase in the rate of cell passage through the cycle and an increase in cell proliferation. These facts indicate a violation of the mechanisms of cell cycle regulation as a result of the action of ionizing radiation (Can and Hicks 2006, Deckbar et al. 2011).

The endocycle is an alternative to the mitotic version of the cell cycle. It is also called the cell cycle of terminal differentiation (Larkins et al. 2001). The consequence of successive cycles of endoreduplication is the formation of polytene chromosomes in the cell nuclei. The polyteny deserves attention as one of the effective mechanisms for enhancing gene expression in eukaryotes. In the literature, various aspects of the adaptive and evolutionary significance of this phenomenon are discussed (Nagl 1976, Edgar and Orr-weaver, 2001, Lee et al. 2009).

One of the consequences of irradiation in the offspring of exposed parents is an increase in the level of embryonic mortality, the so-called dominant lethal mutations. At the population level, this leads to changes in its genetic structure. In turn it affects the general fitness of individuals and its components (Izmailov et al. 1993, Golub and Chernyk 2008). In connection with this, it is of interest to study the features of the functioning of the genome in the progeny of irradiated organisms. Important questions are: (1) whether the effects of ionizing radiation persist in the next generation? and (2) in what way biological systems (organisms, populations) overcome the effects of radiation damage in subsequent generations after the exposure?

The purpose of this investigation was to study effect of single-entry acute γ-irradiation of parent adults on the endoreduplication of giant chromosomes in F1 generation of *Drosophila melanogaster* Meig. The tasks were to investigate the dependence of the effects on irradiation dose, sex, and to determine the statistical power of the influence of these factors on the degree of chromosome polyteny in the progeny of flies.

## Materials and methods

### Biological material and environmental conditions

As a material used wild-type strain *Oregon-R* of *Drosophila melanogaster* Meig. from the collection of the Department of Genetics and Cytology of VN Karazin Kharkiv National University. Flies were grown on a standard sugar-yeast nutrient medium at a temperature of 24.0 ± 0.5°C. *Drosophila* cultures developed in 60 ml vials with 10 ml of the culture medium. Flies laid eggs for 5 days. Larvae for the experiment were taken in the first two days of emergence on the glass.

### Exposure to γ-irradiation

Doses of 8 Gy, 16 Gy and 25 Gy were used in the experiments. Virgin females and males of *Drosophila* adults at the age of 3 days were irradiated with a linear electron accelerator LEA-10. Irradiation was carried out by bremsstrahlung γ-quanta, formed during the interaction of an electron beam with a thick aluminum target. The electron energy was 9.4 MeV, the current strength was 810 μA, the thickness of the aluminum converter was 38 mm. The dose rate at the irradiation point was calculated using Harwell Red 4034 detectors (Harwell, UK), and was 0.4 Gy/sec. The brake spectrum, taking into account the geometry of the experiment, was calculated using the GEANT 4 software package. The brake spectrum was the Bethe-Heitler curve, where 97% of the γ-ray energy was up to 3 MeV, including 70% of energy up to 500 KeV.

### Determination of polyteny degree of chromosomes

The polytene chromosomes were studied in the squashed preparations of *Drosophila* salivary glands, stained with acetoorsein: 2% orcein (Merck KGaA, Darmstadt, Germany) in 45% acetic acid solution (Reahimtrans, Kyiv, Ukraine). The preparations were obtained at the stage of the wandering larva in the late 3rd instar.

Giant chromosomes were examined with a light microscope (MBI-6, “LOMO”, St. Peterburg, Russia). Differences in polyteny degree were determined by the cytomorphometric method (Strashnyuk et al. 1995). Control measurements of the width of chromosomes were carried out in the region of disk 22A of chromosome 2L at 600 × magnification. The ratio of the classes of nuclei with different polyteny degree was studied at 200 × magnification.

We investigated the distribution of nuclei with different level of polyteny in the total preparations of the salivary glands. Based on these data, we calculated the average polyteny degree of chromosomes in normal conditions and after exposure to γ-irradiation.

### Determination of developmental rate

To assess the dynamics of endoreduplication in ontogenesis, we correlated the degree of polyteny in F1 offspring after irradiation with the rate of flies development. The rate of development was studied in synchronized cultures of *Drosophila*. Four-days old *Drosophila* females after mating with the males laid eggs for three hours. In each vial, 20 females were placed. The number of adults released was counted every 3 hours from the beginning to the end of their exit. Males and females were accounted for separately. The results present the combined data of three replicates of the experiment, which showed a high degree of coincidence.

### Statistical methods

Statistical analysis of the experimental data was carried out. The data are presented as the mean ± standard error. Ten larvae in each variant of the experiment were studied. On average, 148–213 nuclei per preparation were studied. In total, from 1484 to 2134 nuclei were analyzed in each experimental group.

The verification of data distributions for compliance with the normal law was carried out using the Shapiro-Wilk test. The significance of the differences in the distribution of nuclei with different polyteny degree of chromosomes was determined by the chi-square test. To distinguish differences in the average degree of polyteny, a two-factor analysis of variance was used with the assessment of statistical impact power of the radiation exposure and the sex. Multiple comparisons were made using the Tukey-Kramer test and Dunnett’s test.

For the analysis of the point parameters of the development rate, the criterion χ2 was used. As point estimates, we used the median time of development. To compare the distributions in different variants, the Kraskell-Wallace test was used, followed by multiple comparison with the control, using the Dunn test.

Differences were considered valid at *p* < 0.05.

## Results

According to Rodman (1967), the initiation of new cycles of endoreduplication in polytene chromosomes stops a few hours before larval-prepupal molt. The giant chromosomes reach at this time the polyteny degrees 256C, 512C, 1024C, and 2048C.

Each cycle of endoreduplication results in a two-fold increase in the number of chromatids in polytene chromosomes. Therefore, nuclei with different levels of polyteny can be easily distinguished visually. In cytological preparations, chromosomes with different polyteny degree differ in width and intensity of staining (Kiknadze and Gruzdev 1970, Strashnyuk et al. 1995). The thickness of chromosomes of different classes of nuclei in the region of the 22A disk used for control measurements was 1.6, 2.3, 3.2, and 4.6 μm. Chromosomes with greater polyteny more intensely stained with acetoorsein.

The polyteny degree of chromosomes (PDC) varies in different parts of the salivary gland: in the distal part it is higher than in the proximal (Figure 1). The correspondence between the cytomorphometric characteristics of polytene chromosomes and their degree of polyteny was demonstrated earlier (Strashnyuk et al. 1995, Dyka et al. 2016). The number of classes of nuclei with different width of the chromosomes, their location in the gland and percentage showed the close compliance with Rodman’s (1967) cytophotometry data.

**Figure 1.**
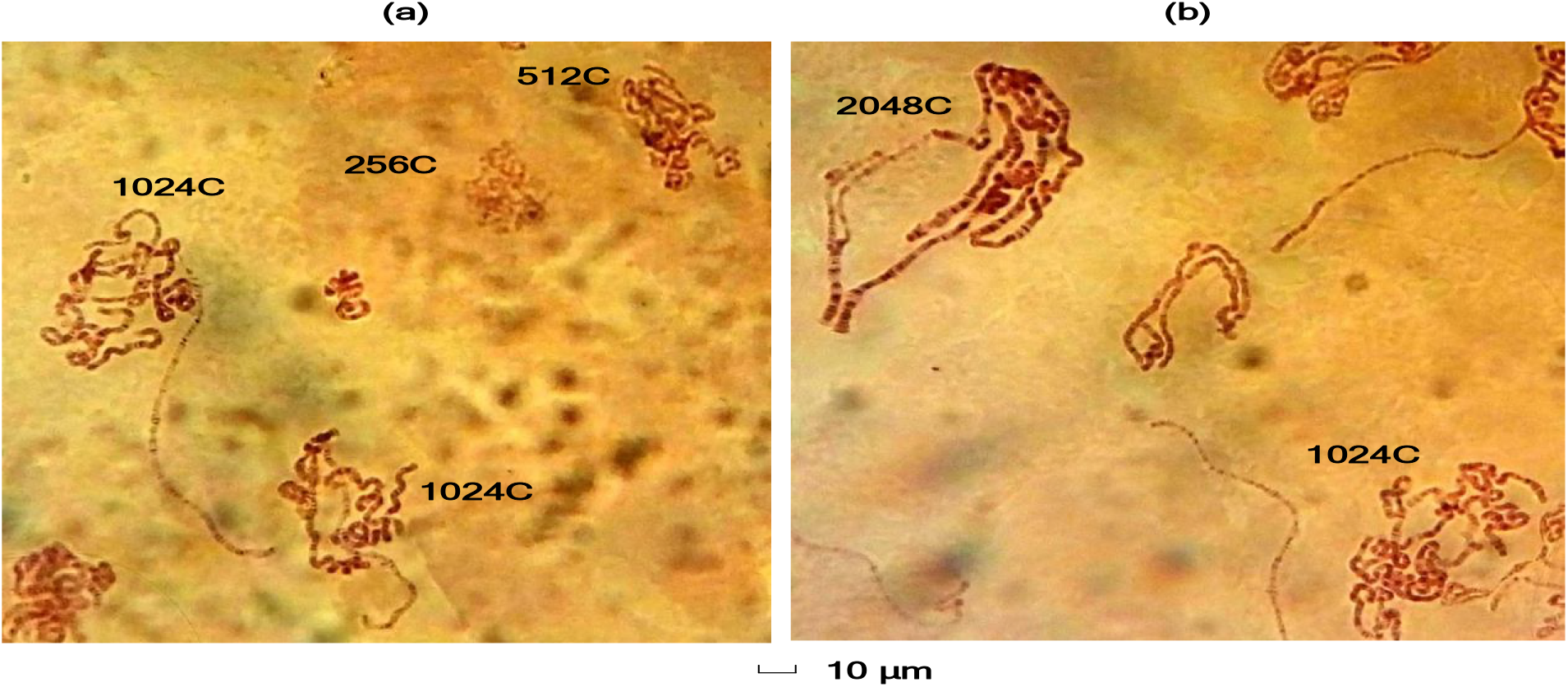
Giant chromosomes of *Drosophila melanogaster* stained by acetoorcein with different polyteny degree: (a) proximal part of the salivary gland; (b) distal part of the salivary gland.

Figure 2 presents data on the distribution of nuclei with different polyteny degree of chromosomes in the salivary glands of *Drosophila* larvae in the F1 generation after γ-irradiation. In males, irradiation at a dose of 8 Gy caused a decrease in the fraction of 256C and 512C nuclei and an increase in the percentage of 1024C and 2048C nuclei. A similar effect occurred at a dose of 25 Gy, with the exception of the fraction of 256C nuclei that did not differ from the control values. At a dose of 16 Gy, on the contrary, an increase in the percentage of nuclei 256C and 516C was observed, while the percentage of 1024C nuclei was lower than in the control.

**Figure 2.**
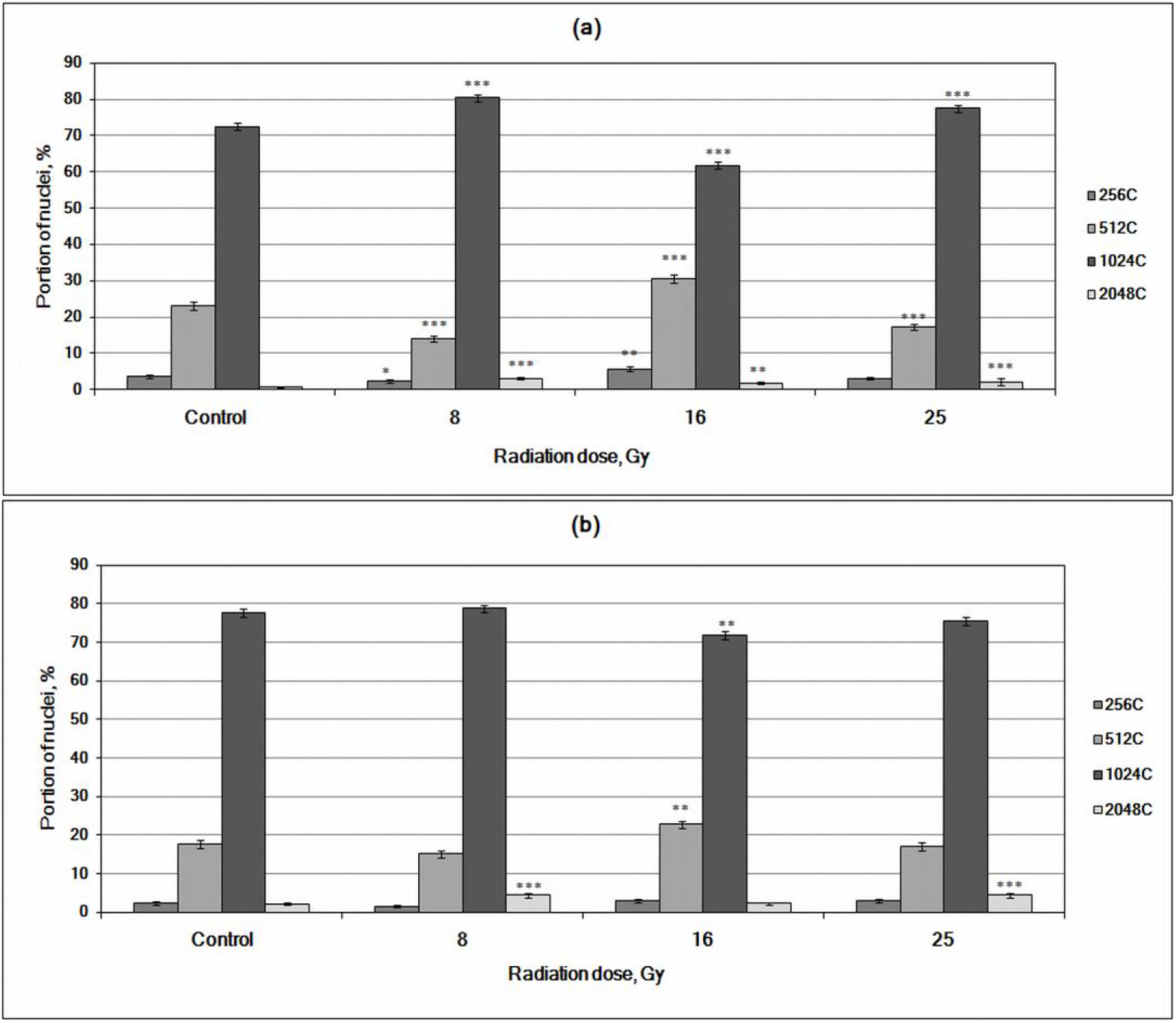
The distribution of nuclei with different polyteny degree in *Drosophila melanogaster* salivary glands in F1 generation after γ-irradiation: (a) males; (b) females. * *p* < 0.05; ** *p* < 0.01; *** *p* < 0.001: versus to control group.

In females, the changes were less significant. At the dose of 8 Gy the portion of 2048C nuclei increased slightly. At the dose of 16 Gy, the content of 512C nuclei was higher, and the number of 1024C nuclei decreased. At the dose of 25 Gy, the distribution of nuclei with different degrees of polyteny did not differ from the control values.

Data on the percentage of nuclei with different genome ploidy was used for calculation of averages of polyteny degree in the salivary glands of *Drosophila* larvae in the control and in experimental variants. The results are shown in Figure 3.

**Figure 3.**
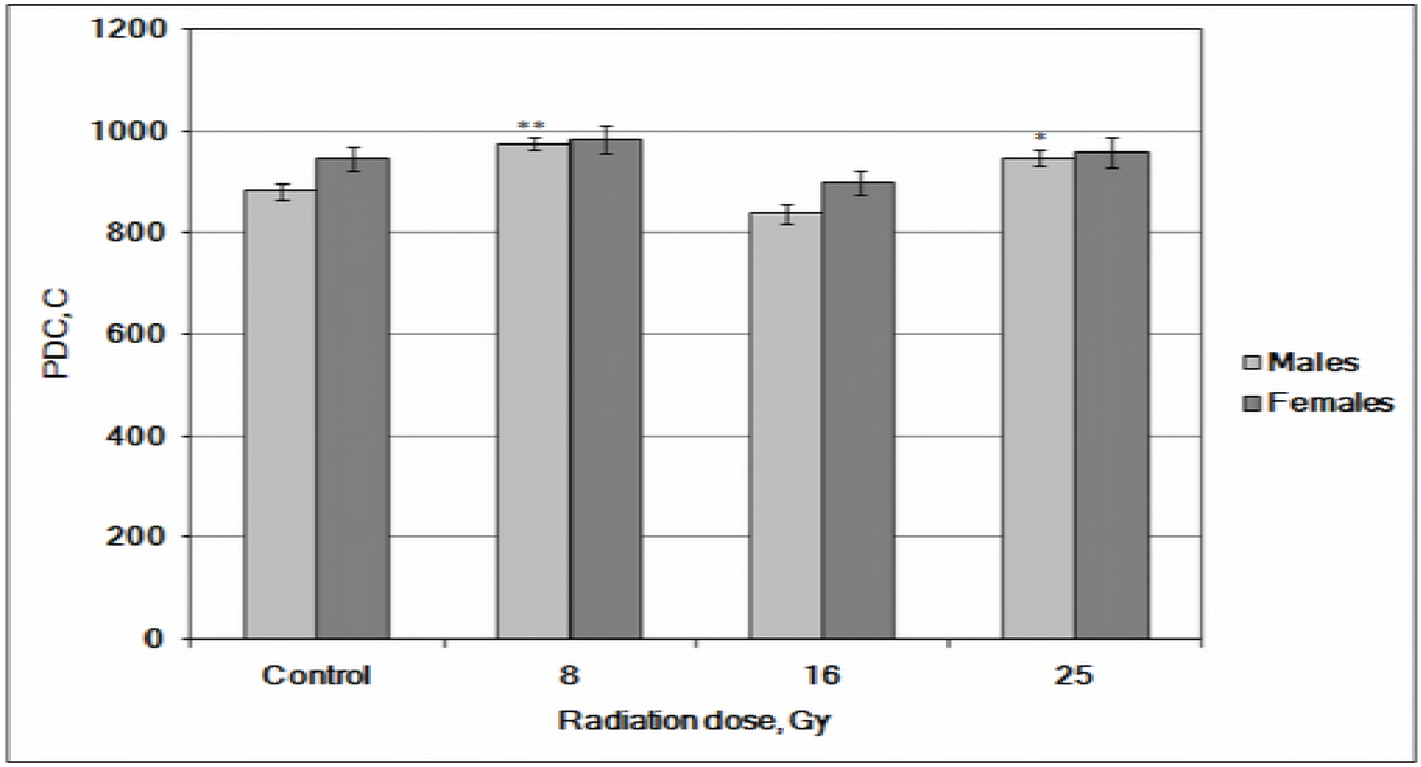
The average values of polyteny degree of chromosomes (PDC) in *Drosophila melanogaster* salivary glands in F1 generation after γ-irradiation. * *p* < 0.05; ** *p* < 0.01: versus to control group.

In males, the mean values of polyteny in F1 generation after γ-irradiation were higher than the control values at the dose of 8 Gy by 10.6%, and at the dose of 25 Gy – by 7.4%. At the dose of 16 Gy, the average PDC in males did not show significant changes. This means that changes in the distribution of nuclei with different polyteny at this dose were compensatory in nature.

In females, the mean values of polyteny degree of chromosomes after γ-irradiation of parents did not differ from the control, irrespective of the irradiation dose.

The data presented in Figure 4 show that the rate of development did not change after irradiation in dose of 8 Gy. The development significantly accelerated at 25 Gy: the median decreased by 5.3 hours (*p* < 0.001). Thus, the increase in the degree of polyteny in males at a dose of 8 Gy and even more at 25 Gy is a consequence of an increase in the level of endoreduplication and is not associated with an elongation of the developmental period. Implicitly, these data also demonstrate a dose-response effect on endoreduplication.

**Figure 4.**
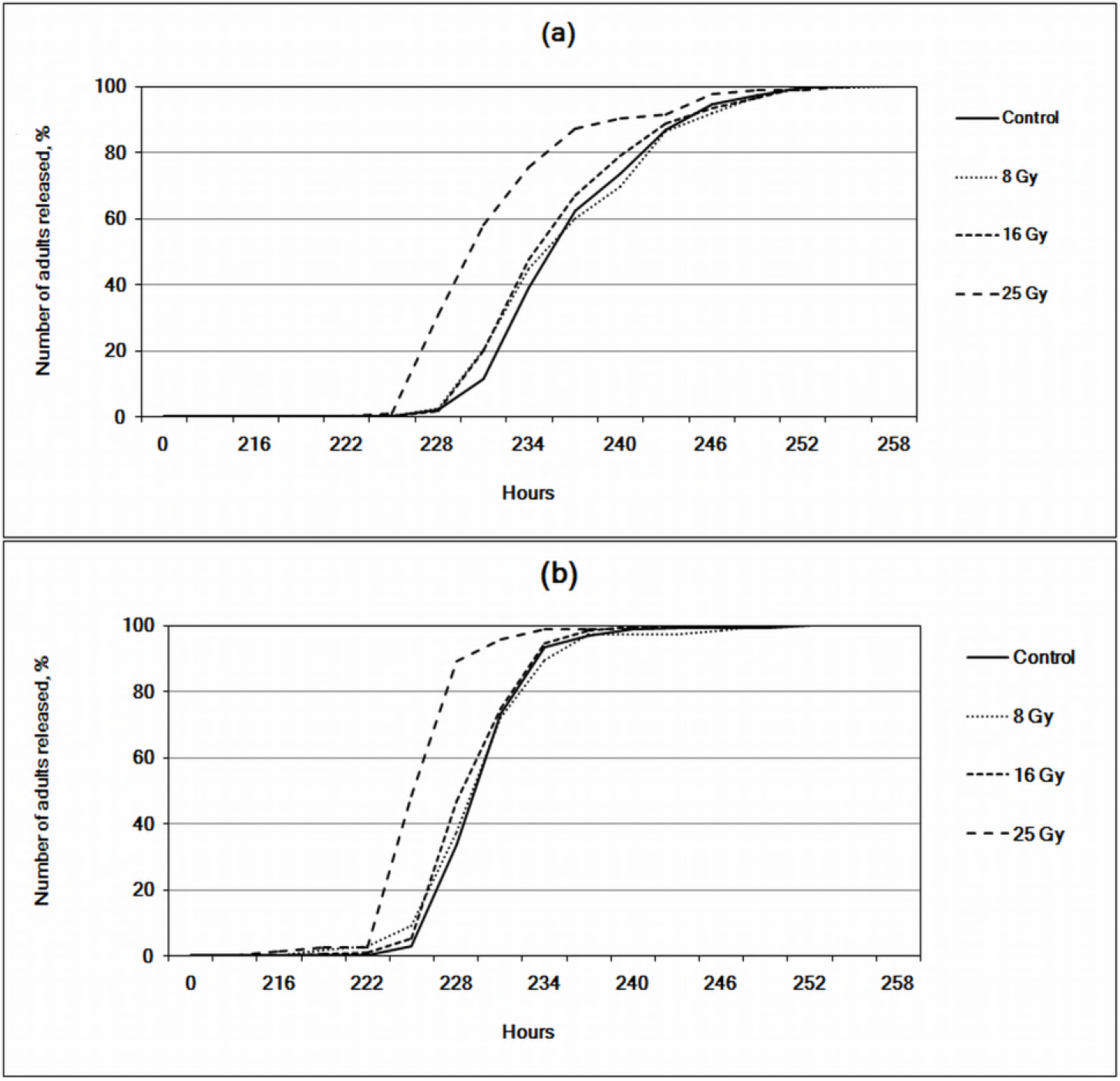
The rate of development in *Drosophila melanogaster* in F1 generation after acute γ-irradiation: (a) males; (b) females.

In females, development was also significantly accelerated after irradiation at a dose of 25 Gy. The median decreased by 4.1 hours (*p* < 0.001). Some acceleration of development, although less significant, was also observed at a dose of 16 Gy in males. The median decreased by 1.0 hour (*p* < 0.05). The final polyteny in these cases did not differ from the control values. However, this result was achieved in a shorter time. Consequently, endoreduplication also occurred more actively.

The obtained data indicate that the degree of genome amplification in the salivary glands of *Drosophila* after γ-irradiation of the parental individuals depends on two factors: sex and radiation dose. To estimate the statistical impact power of the studied factors on endoreduplication the variance analysis of two-factor complexes was used. The impact power of the factor is defined as the fraction of factorial variability in the overall variability of the trait. The results of the analysis are presented in the Table 1. According to the data obtained, the statistical impact power (*ή^2^*) of the sex on the polyteny degree of chromosomes was 4,9%, the radiation impact was 26,8%. The combined effect of the two factors was not significant.

**Table 1.**
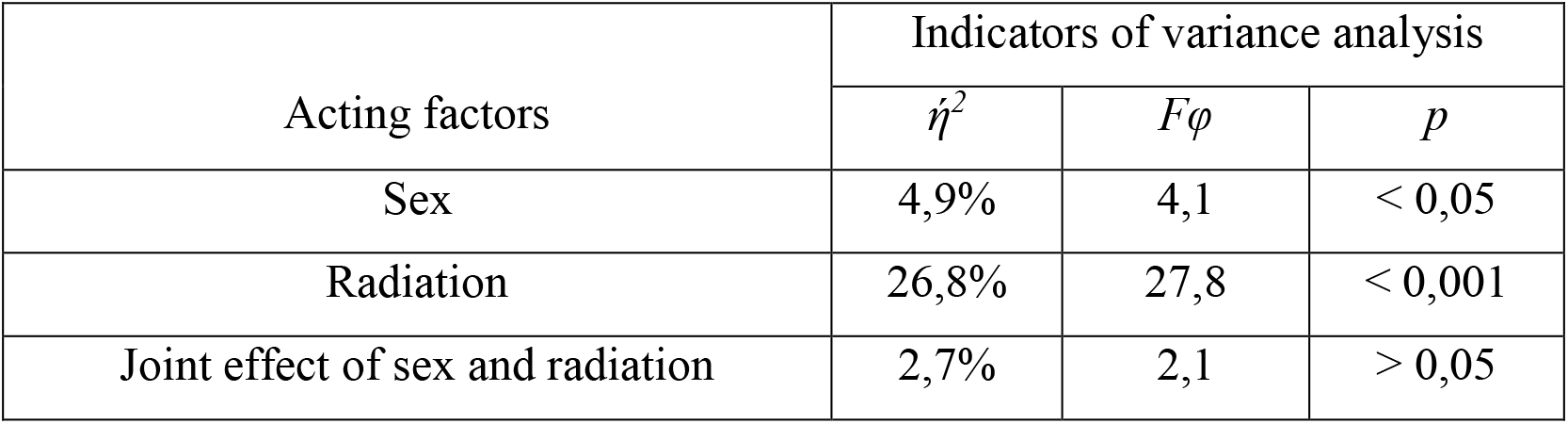
The statistical impact power of radiation and sex on the polyteny degree of chromosomes in F1 generation of *Drosophila melanogaster*.

## Discussion

Genome amplification by endoreduplication is a characteristic phenomenon for cells of many differentiating tissues of eukaryotes. Endoreduplication is an effective mechanism for enhancing gene expression and increasing the metabolic potential of cells. Endocycles also promote accelerated growth (Marguerat and Bähler 2012), response to physiological stress (Zhuravleva et al. 2004, Fox and Duronio 2013) and adaptation to environmental conditions (Strashnyuk et al. 1997, Zhuravleva et al. 2004). According to experts (Sugimoto-Shirasu and Roberts 2003, Zielke et al. 2011), endoduplication produces about half of the world biomass. This indicates the biological significance of this phenomenon.

At the cellular level, the endocycle is controlled by key regulators of the cell cycle, such as cyclins, cyclin-dependent kinases and their inhibitors (Larkins et al. 2001, Zielke et al. 2011, Fox and Duronio 2013). Humoral factors also play an important role. In particular, the effects of the juvenile hormone (Sihna and Lakhotia 1983) and ecdysterone (Shakina and Strashnyuk 2011) on the endoreduplication process in *Drosophila* were shown.

A significant contribution to the variability of polyteny degree of chromosomes is made by hereditary factors (Strashnyuk et al. 1995, Larkins et al. 2001). In addition, the modifying effect on the ploidy of cells is exerted by external conditions, such as temperature (Strashnuk et al. 1997), culture density (Rarog et al. 1999, Zhuravleva et al. 2004), food composition (Britton and Edgar 1998).

As for present study, we believe that the observed changes in the chromosomes polyteny degree in the progeny of irradiated parents were due to the selection factor. Embryonic mortality in the *Oregon-R* strain after irradiation of adults in the dose range of 8–25 Gy was 2.1–4.3 times higher than the control values. The effect was dose-dependent. Gametic selection could also contribute to the variability of the trait after ionizing irradiation (Hourcade et al. 2010).

Earlier we showed changes in the offspring fitness after irradiation of the parents in *Drosophila*. The lifespan of adults in F1 increased or did not change (Skorobagatko et al. 2016). Under conditions analogous to our experiment, at a dose of 25 Gy, the average lifespan increased in males, in females it did not change. Izmailov et al. (1993) also observed increased longevity of flies in the first generation after irradiation of parents.

The frequency of dominant lethal mutations increased in F1 progeny after irradiation, but returned to the control values or (at 25 Gy) decreased in the progeny of F2 (Skorobagatko et al. 2015). This indicates the appearance of genetic changes in the strain, at least at a dose of 25 Gy. Thus, selection did occur, and the offspring after that became more viable.

We can also assume the effect of hormesis, that is, the action of epigenetic mechanisms. Epigenetic mechanisms begin to act already when the egg is formed, when gradients of concentrations of biologically active substances are formed (Korochkin 2006). We applied the exposure to radiation at this stage. However, the hormesis effect requires justification (Mushak 2007). In our case, this is difficult to do, since the selection takes place. If we are talking about epigenetic phenomena, then we must take into account that they do not concern the changes in the genotype. At the same time, the epigenetic mechanisms of the action of radiation are discussed in the literature (Vaiserman et al. 2004, Sarup and Loeschcke 2011). Perhaps the different mechanisms operate at different doses.

The differences between males and females in response to the action of radiation may be explained by different viability of the sexes. It is known that the homogametic sex in this respect is superior to the heterogametic sex. This follows from the well-known Haldane rule (Haldane 1922), as well as the hypothesis of sex-linked lethal and semi-lethal genes (Huxley 1924).

Geodakyan (1998) considers the phenomenon of sexual differentiation from the standpoint of their specialization at the population-species level. According to his view, evolutionary innovations in the male genome occur before they are transferred to the female genome. This can be explained from the positions of dichronic evolution, when the evolutionary changes in the males are faster than in the females.

In our study, the changes in polyteny are detected in males, and in females they are absent. The polyteny in males increased. This suggests that selection for an increase in radioresistance implies an increase in the metabolic potential of cells.

Some authors suggest as a possible function of polyteny the modulation of stress response (Cookson et al. 2006) or buffering of genome (Edgar and Orr-weaver 2001). In our previous works we found that the differences in polyteny degree of chromosomes in *Drosophila* positively correlated with heat resistance, body weight of adults, and general fitness (Strashnyuk et al. 1995, 1997, Zhuravleva et al. 2004). According to Hassel et al. (2014), cells in which the endocycle occurs are less likely to respond to DNA damage, for example, in the case of radiation-induced instability of the genome.

In response to the effect of ionizing radiation, the E2F1 transcription factor is overexpressed (Wichmann et al. 2010). E2F1 is the central component of the endocyclic moleculal oscillator which regulates the periodic expression of Cyclin E. In turn, Cyclin E catalyzes kinase CDK2 in the G-S transition (Zielke et al. 2011).

Endocycles also contribute to the repair of damaged tissues, which is an alternative or complement to the function of stem cells (Losick et al. 2013, Xiang et al. 2017).

The above facts can be useful for explaining the possible connection between the selection for an increase in the radioresistance and the degree of genome amplification in *Drosophila* polytene chromosomes after exposure to γ-irradiation.

## Conclusions

Endoreduplication in *Drosophila* F1 generation after acute γ-irradiation of the parental adults increases. The average means of polyteny degree of chromosomes in males were higher than the control values at the irradiation dose of 8 Gy by 10.6%, at the dose of 25 Gy by 7.4%, and at the dose of 16 Gy did not show significant changes. In females, the average polyteny did not differ from the control value irrespective of the irradiation dose. Such polyteny values in the offspring of irradiated flies were achieved at a faster rate of development. The statistical impact power of sex on the chromosomes polyteny degree was 4,9%, the radiation impact was 26,8%. The enhancement of endoreduplication is considered as a consequence of an increasing selection pressure after irradiation.

## Declaration of interests

The authors report no conflicts of interest. The authors alone are responsible for the content and writing of the paper.

## Funding information

This work was supported by the Ministry of Education and Science of Ukraine (Project State registration number: 0117U004836).

